# Benchmarking of protein interaction databases for integration with manually reconstructed signaling network models

**DOI:** 10.1101/2022.10.25.513640

**Authors:** Matthew W. Van de Graaf, Taylor G. Eggertsen, Angela C. Zeigler, Philip M. Tan, Jeffrey J. Saucerman

## Abstract

Protein interaction databases are critical resources for network bioinformatics and integrating molecular experimental data. Interaction databases may also enable construction of predictive computational models of biological networks, although their fidelity for this purpose is not clear. Here, we benchmark protein interaction databases X2K, Reactome, Pathway Commons, Omnipath, and Signor for their ability to recover manually curated edges from three logic-based network models of cardiac hypertrophy, mechano-signaling, and fibrosis. Pathway Commons performed best at recovering interactions from manually reconstructed hypertrophy (137 of 193 interactions, 71%), mechano-signaling (85 of 125 interactions, 68%), and fibroblast networks (98 of 142 interactions, 69%). While protein interaction databases successfully recovered central, well-conserved pathways, they performed worse at recovering tissue-specific and transcriptional regulation. This highlights a knowledge gap where manual curation is critical. Finally, we tested the ability of Signor and Pathway Commons to identify new edges that improve model predictions, revealing important roles of PKC autophosphorylation and CaMKII phosphorylation of CREB in cardiomyocyte hypertrophy. This study provides a platform for benchmarking protein interaction databases for their utility in network model construction, as well as providing new insights into cardiac hypertrophy signaling.

## Introduction

Systems modeling aims to quantitatively predict molecular signaling and provide insight into the complex dynamics of biochemical processes. A number of groups have developed manually curated signaling networks and used the network reconstructions to produce quantitative, mechanistic models (1–3). One approach employs a logic-based differential equation modeling approach for cell signaling networks based on normalized Hill activation functions controlled by AND and OR gates to characterize signaling interactions. Because this approach uses default parameters to predict fractional activation of network species, the modeling technique can be employed directly with data on directed interactions. Using this approach, we have developed network reconstructions for cardiac myocyte hypertrophy signaling [2], cardiac fibroblast differentiation signaling [3], and cardiomyocyte mechano-signaling [4]. While each of these models has demonstrated accuracy when validated against published literature, the process of developing the networks requires labor and time-intensive manual curation. More extensive use of protein interaction networks may enable more systematic model building. However, protein interaction networks contain gaps and false positives (7), so it is unclear how useful they may be.

In this study, we used three manually curated network models of cardiac myocyte hypertrophy signaling [2], cardiac fibroblast differentiation signaling [3], and cardiomyocyte mechano-signaling [4] to benchmark the quality of popular protein interaction databases X2K [5], Signor [6], Reactome [7], Omnipath (11) and Pathway Commons [8]. Our benchmarking focused on evaluating the quality of undirected and directed interactions that would be useful for constructing predictive network models. We also examined what regions of a network have better coverage by interaction databases. Finally, Pathway Commons and Signor were used to guide expansion of the hypertrophy signaling network and provide new insights into hypertrophy.

## Results and Discussion

### Benchmarking against the cardiac hypertrophy signaling network

In order to benchmark protein interaction databases against manually curated network reconstructions, it is necessary to first annotate the genes or genes that correspond to each node in the network model. A single node may represent multiple protein isoforms or a protein complex constituted of subunits. The manual curation process of constructing these networks requires deciding which proteins, and consequently which genes, constitute a given node based on the available information from literature. For example, node A is listed to represent genes A1 and A2 in the workflow schematic (**Figure 1**). In benchmarking any reactions containing node A against a database, both genes A1 and A2 would be considered possible representations of the node. Using this approach, we translated the model’s reactions into gene-gene edges.

**Figure 1.**
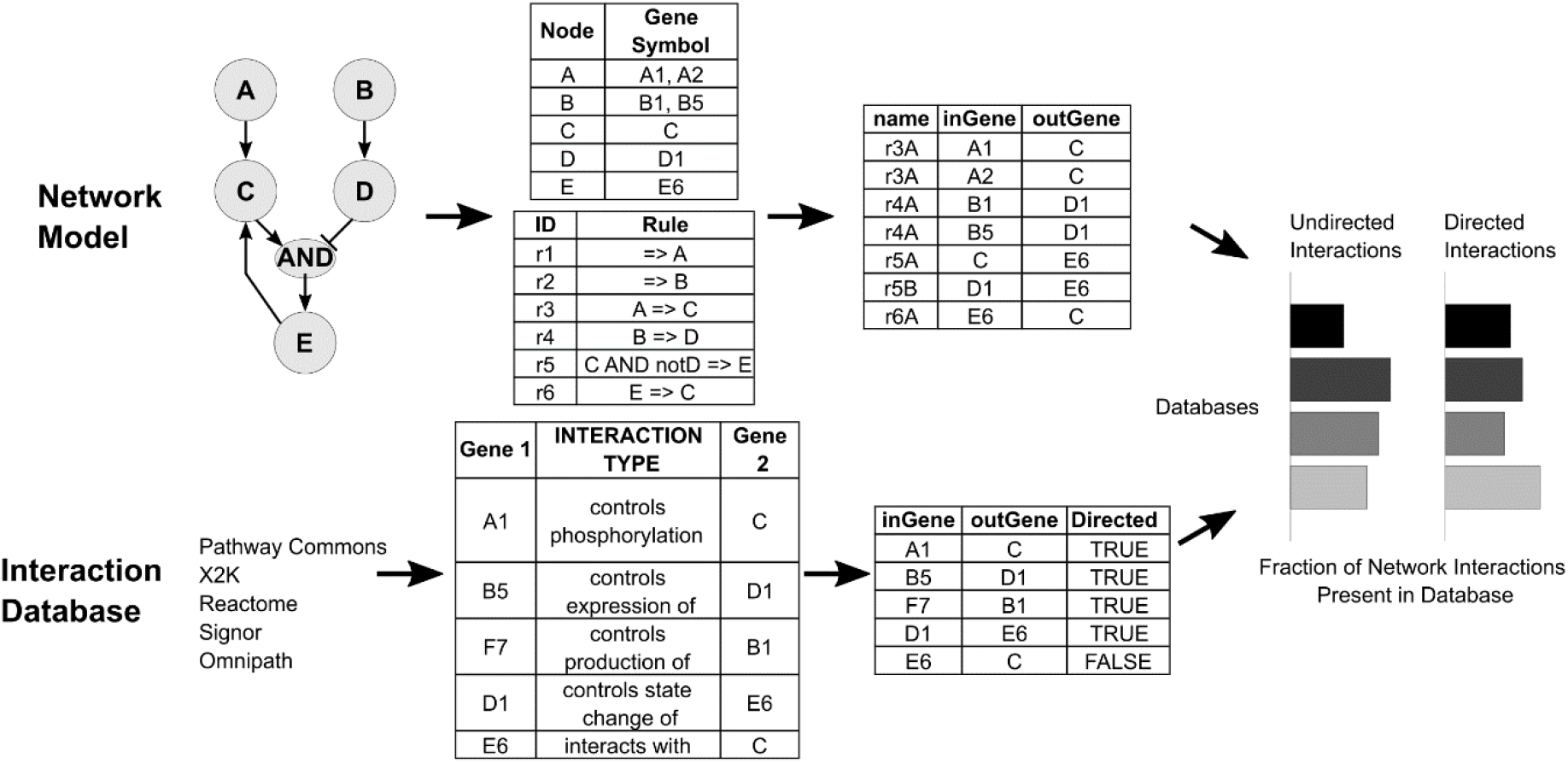
Process used to benchmark protein interaction databases against manually curated network models. Manual network reconstructions were translated into a tabular format that matched files obtained from protein interaction databases.

Among the largest and most popular protein-interaction databases, X2K [5], Signor [6], Reactome [7], Omnipath (11) and Pathway Commons [8] were selected for benchmarking. The available datasets for each of these sources were downloaded for use in analysis. For X2K, it was necessary to first concatenate the individual files provided for data sources online. In each of the databases used for benchmarking, the type of interaction is denoted by a separate field. In the example database (**Figure 1**), the top four interactions are directional, whereas the fifth interaction is undirected. As shown in the figure, the toy network would be compared to all five reactions to produce a benchmarking score. Benchmarking scores are computed for just undirected interactions, just directed interactions, and for all interactions in the specified database.

We next applied this benchmarking framework to benchmark five protein interaction databases in their ability to recover edges from a logic-based differential equation model of cardiomyocyte hypertrophy signaling composed of 193 manually curated edges. Benchmarking X2K [5], Signor [6], Reactome [7], Omnipath (11) and Pathway Commons [8] for undirected interactions, directed interactions, and all interactions revealed variability in the overall content of specific databases (**Figure 2b**). For example, the X2K database performs well when only considering undirected interactions but poorly for directed interactions. However, it should be noted that directed interactions are critical for model construction. In contrast, Signor and Omnipath perform well for directed interactions but have little information for undirected interactions. For the directed, undirected, and combined benchmarking analysis, Pathway Commons outperformed all other databases. Pathway Commons exhibited a reasonably good benchmarking score for directed (62%), undirected (57%), and all (directed and undirected) interactions (71%). Notably, these results represent the expected results based on the size and content of each database (**Table 1**). Pathway Commons contains the largest number of interactions, while X2K contains mostly undirected interactions and Signor and Omnipath perform well despite having much fewer directed interactions than Pathway Commons. We next asked whether combining interaction databases would further improve benchmarking. The union of all five databases benchmarked only marginally higher than Pathway Commons.

**Figure 2.**
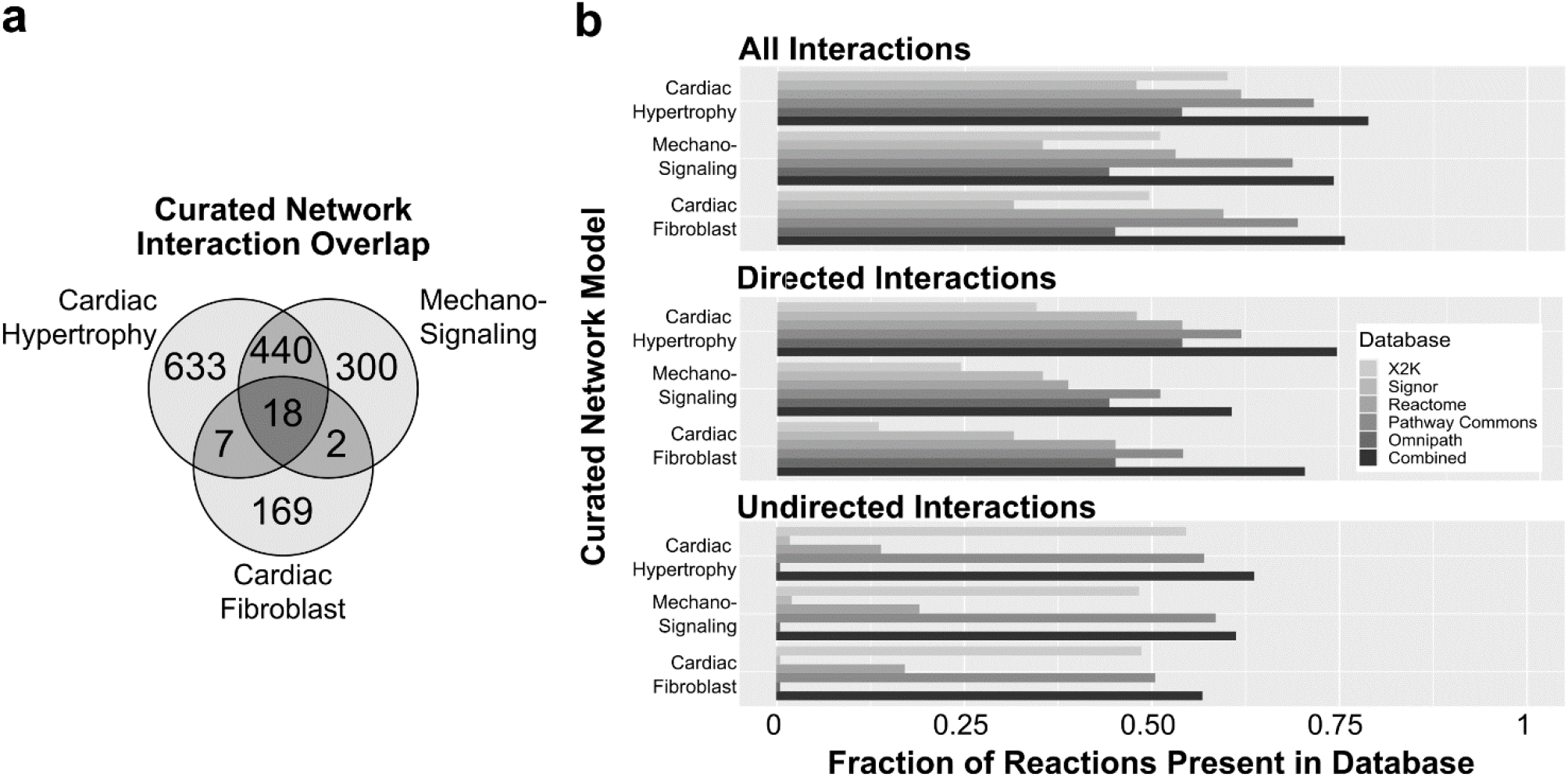
Benchmarking of five protein interaction databases against three manually reconstructed network models. a) Overlap in reactions between the three manually curated network models (hypertrophy, mechano-signaling, fibroblast). Network edges largely retain distinct biological interactions when converted to gene product interactions. b) Manually curated network models of cardiac hypertrophy, mechano-signaling, and fibroblasts were used to benchmark to five biological interaction databases (X2K, Signor, Reactome, Pathway Commons, Omnipath) and their combination. In each case, we quantified benchmarking scores from the fraction of reactions from the manual network that could be recovered from the interaction database. Undirected and directed interactions could be merged to produce an overall recovery score.

**Table 1.** Quantity and type of interaction information in databases.

| Database Name | Directed Interactions | Undirected Interactions | Total |
| --- | --- | --- | --- |
| Expression2Kinases (X2K) | 11,549 | 318,485 | 330,034 |
| Pathway Commons | 479,298 | 508,480 | 987,778 |
| Reactome | 99,135 | 131,108 | 230,243 |
| Signor | 18,112 | 1,407 | 19,519 |
| Omnipath | 40,014 | 0 | 40,014 |

### Extending analysis to fibroblast and mechano-transduction signaling networks

The cardiac hypertrophy signaling network [2], cardiac fibroblast differentiation network [3], and the cardiac mechano-signaling network [4] each represent large-scale network reconstructions of similar size with respect to the number of nodes and edges included (**Table 2**). Across the three networks used for benchmarking, there is an inevitable level of overlap due to common signaling pathways having functions across cell types and diseases. Prior to benchmarking, the uniqueness of the three networks was quantified by measuring overlap in the gene product interactions represented (**Figure 2a**). Despite the presence of overlap in gene products, each network contains a large number of distinct reactions. Thus, expanding the benchmarking analysis to two additional manually reconstructed signaling networks demonstrates the ability to generalize benchmarking across other networks and models.

**Table 2.** Number of nodes, genes, and edges represented in network models.

| Network | Nodes | Genes | Edges |
| --- | --- | --- | --- |
| Cardiac Fibroblast | 91 | 95 | 142 |
| Cardiac Hypertrophy Signaling Network | 106 | 217 | 193 |
| Cardiac Mechano-Signaling | 94 | 229 | 125 |

Benchmarking the mechano-signaling and cardiac fibroblast signaling networks against the five databases largely mirror that of the cardiac hypertrophy signaling network (**Figure 2b**). X2K and Signor predominantly contain undirected and directed information, respectively, and Pathway Commons performed the best across all three classes of comparison. Like benchmarking against the hypertrophy network, the combination of directed and undirected interactions from Pathway Commons was able to recall 68% of the mechano-signaling network and 69% of the fibroblast network. Consistent with results from the cardiac hypertrophy signaling network, Signor demonstrates valuable performance (35% of mechano-signaling, 31% of fibroblast) with regard to directed interactions despite having only 3.8% as many directed interactions as Pathway Commons. Omnipath has the greatest specificity of directed interactions, and exhibits high performance for a moderate sized database (8.3% as many directed interactions as Pathway Commons).

### Benchmarking protein interactions to networks using Pathway Commons interaction annotations

Protein interaction databases provide not only the interactions themselves but also annotations on the type of interactions. We asked whether certain types of interaction annotations were more associated with successful benchmarking to edges in the three network models. Within the Pathway Commons interaction database, there are seven different interaction types used to annotate interactions between proteins. We classified these interaction types as either “functional” interactions, which denote a directed pathway effect of one gene product on the other, or “physical” interactions, which denote a direct chemical interaction between the two gene products.

For each network model used in benchmarking, the fraction of all functional interactions and all physical interactions of each subtype was tabulated and compared to the corresponding values for the entire Pathway Commons database. Curated network models demonstrated enrichment for specific interaction types when compared to the overall Pathway Commons database. For example, looking at functional interactions, the three manually curated networks exhibited a higher percentage of annotations for “controls state change of” and “controls phosphorylation of” compared to the overall Pathway Commons database (**Figure 3a**). In contrast, while the fraction of reactions classified as “catalysis precedes” and “controls expression of” is lower in the manually curated networks. This indicates that specific the process of manually curating a network model may cater to specific types of functional interactions, but it is unclear whether this reflects curation bias or utility. Physical interactions annotated as “interacts with” or “in complex with” were more balanced across manually curated networks and Pathway Commons. The higher proportion of “controls-state-change-of” and “controls-phosphorylation-of” functional annotations among network model edges compared with Pathway Commons suggests that they are particularly valuable for integrating network models with interaction databases.

**Figure 3.**
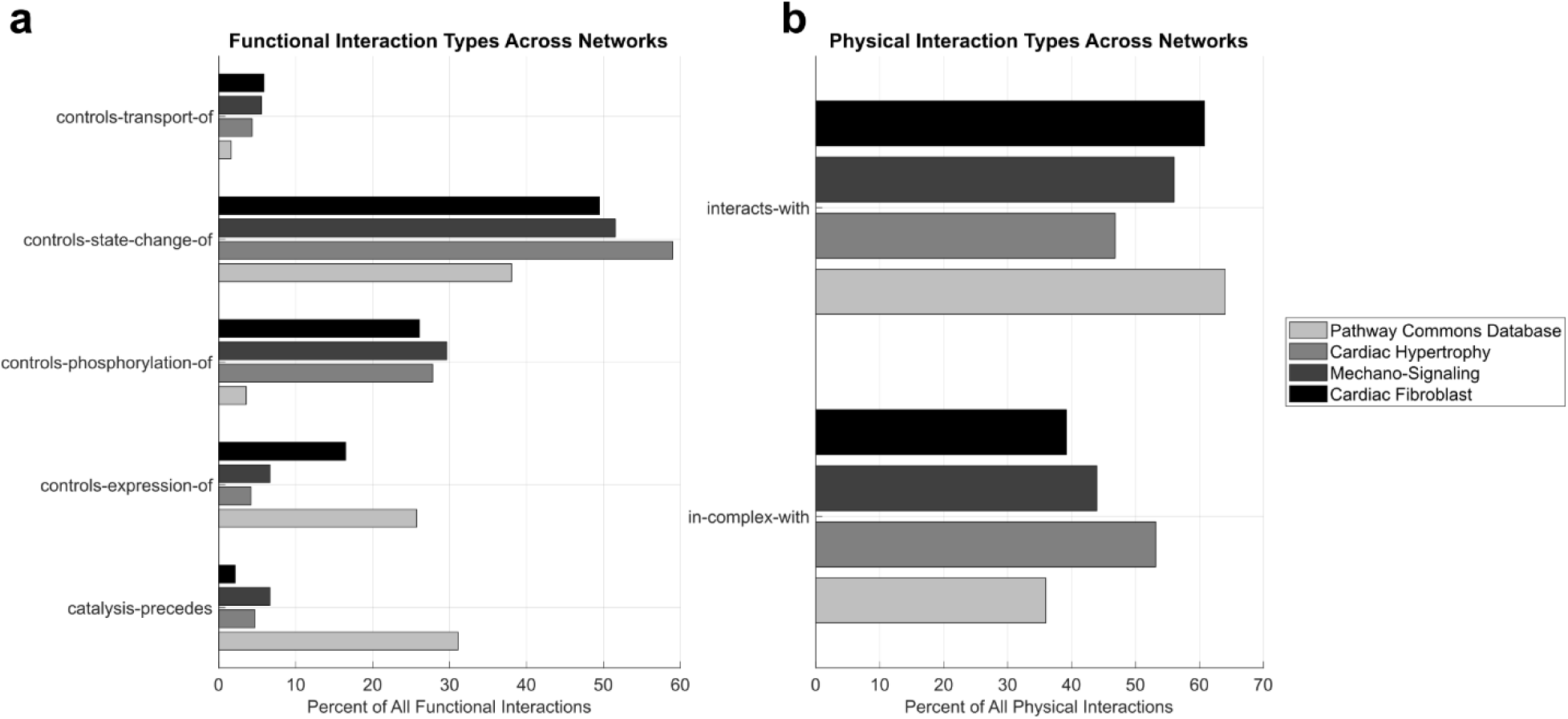
Comparison of interaction types among Pathway Commons compared with cardiac hypertrophy, mechano-signaling, and fibroblast network models. Within the categories of functional and physical interactions, multiple specific interaction descriptors are used to identify the mechanism of the relationship.

We next examined the relative coverage of functional vs. physical interaction annotations from Pathway Commons that mapped to the manually curated hypertrophy, mechano-signaling, and fibrosis networks. The Pathway Commons database itself is composed of a similar balance of functional and physical annotations, with only 1.8% of interactions containing both types of annotation within Pathway Commons (**Figure 4a**). In contrast, all three manually curated networks contained a greater proportion of protein interactions that had support by both functional and physical annotations (31% for hypertrophy, 30% for mechano-signaling, 29% for fibroblast) (**Figure 4a**). This result builds upon the previous interaction annotation subtype analysis in **Figure 3** by demonstrating that individual protein interactions in the curated network models have much greater support in Pathway Commons than seen for typical biochemical interactions.

**Figure 4.**
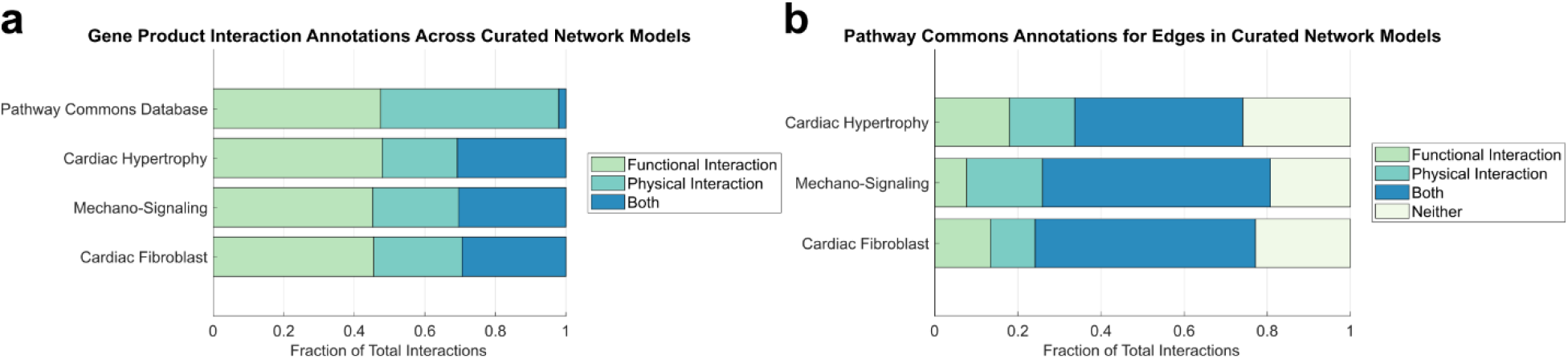
Comparison of annotation support by Pathway Commons annotations for manually curated edges in the hypertrophy, mechano-signaling and fibrosis networks. a) Proportion of interactions from each network model that are annotated by functional or physical annotations from Pathway Commons. b) Merging The results of gene product interaction annotations were then merged into network edges.

The above analysis examined annotation at the level of individual gene or protein interactions, but network edges in the manually curated networks often involve multiple gene/protein isoforms or paralogs. To assess annotation coverage at the network edge level, we merged annotations, essentially reversing the direction of the schematic in **Figure 1**. Due to the multiple protein isoforms or paralogs within each network, the fraction of network edges supported by both functional and physical interactions (40%, 55%, and 52% for hypertrophy, mechano-signaling, and fibroblast networks; **Figure 4b**) is greater than the corresponding values for individual protein interactions (**Figure 4a**). The fraction of network edges supported by either functional or physical Pathway Commons annotation also increased, to 74%, 81%, and 77%, respectively.

### Benchmarking of protein interaction databases across the topology of the hypertrophy network

We next asked whether annotation coverage by interaction databases differ across the topology of a manually curated network, and whether such topology-specific coverage may differ between Pathway Commons (which has the highest overall benchmarking score) and Signor (which has high specificity for directed annotations). As shown in **Figure 5**, network-wide annotations show areas of greater or weaker annotation coverage by Pathway Commons and Signor. In both Pathway Commons and Signor, centrally located network edges are well-known and common across cell types. These edges are annotated by both Pathway Commons and Signor, such as growth factors signaling to receptors. Contrastingly, many cardiac-specific edges or regulation of gene transcription demonstrate less coverage by Pathway Commons or Signor.

**Figure 5.**
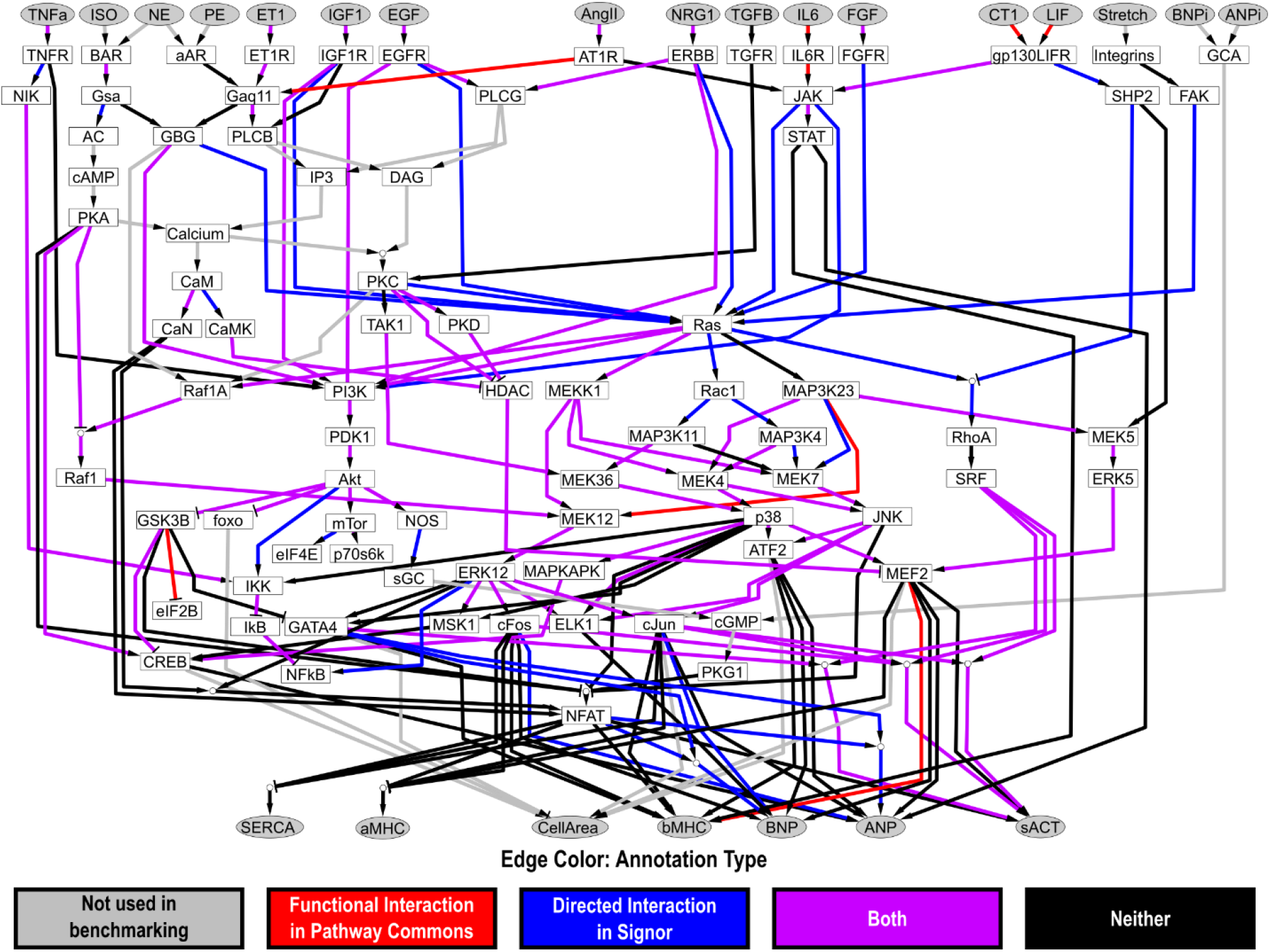
Merged Pathway Commons and Signor annotations visualized on the hypertrophy signaling network. Functional annotations in Pathway Commons and directed annotations in Signor for the cardiomyocyte hypertrophy signaling network demonstrate a large degree of overlap.

### Prioritization of new network edges based on literature support in protein interaction databases

Manual literature curation to expand a network model is challenged by a vast and sometimes inconsistent literature. Now that Pathway Commons and Signor had been benchmarked by the network models, we asked whether these databases could be used to prioritize addition of new network edges that were not captured during manual construction of the hypertrophy network. Initially, we attempted to prioritize candidate edges between proteins already in the network using the number of documented Pathway Commons PubMed IDs for a given edge. However, manual inspection of PubMed IDs supporting one of the top results, Jak => ERK12 revealed that the literature did not support this interaction, suggesting other prioritization strategies were needed. Next, we attempted to infer network edges ranked by the number of PubMed IDs documented by Signor for a given candidate edge. Analysis edges already present in the hypertrophy signaling network (**Table 3**) showed very high support by Signor for MEK36 => p38, Akt => foxo, and FGF => FGFR, which are all well appreciated in the cardiac literature. The Signor-derived metric produced more reasonable results, likely due to the manual curation process being used to create the Signor database.

**Table 3.**
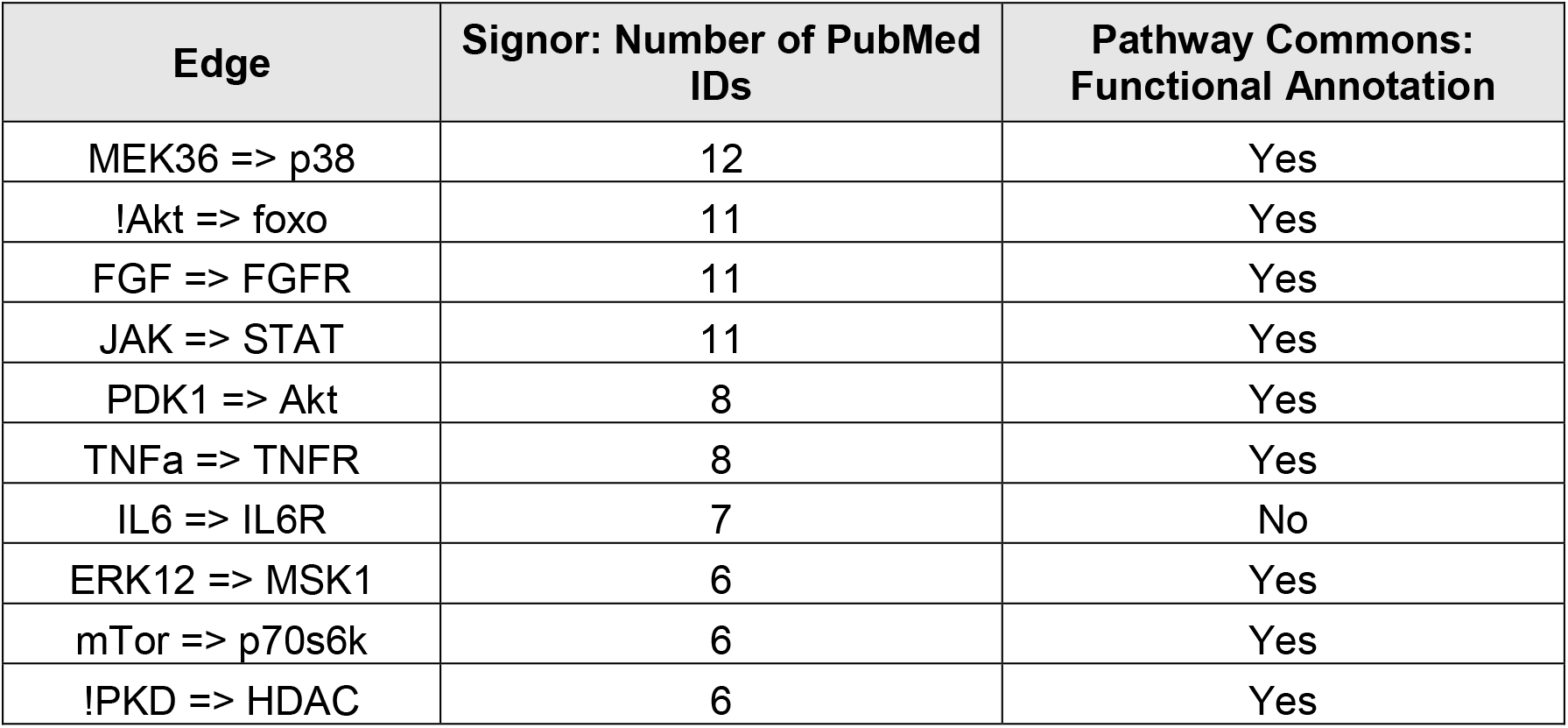
Top 10 network edges among edges already in the hypertrophy network, ranked by number of Signor PubMed IDs.

We next asked whether this database interaction-ranking strategy could identify new candidate edges for inclusion in the network model. As above, we searched Signor for interactions between proteins that were already in the hypertrophy network model, but in this case, we sought interactions that were not in the original hypertrophy network. Notably, seven of the top fifteen candidate edges (**Table 4**) represent auto-phosphorylation interactions that function to sustain activity of a given node. Among the other edges listed in **Table 4**, new possible edges and network refinements are both represented. For example, the ERK12 => MAPKAPK demonstrates a need to more carefully delineate target specificity, as ERK targeted MAPAPKs include ribosomal S6 kinases RSKs, MSKs, and MNKs (13). The PubMed IDs documented for CaMKII => CREB reveal that CaMKII phosphorylates and activates CREB (14).

**Table 4.**
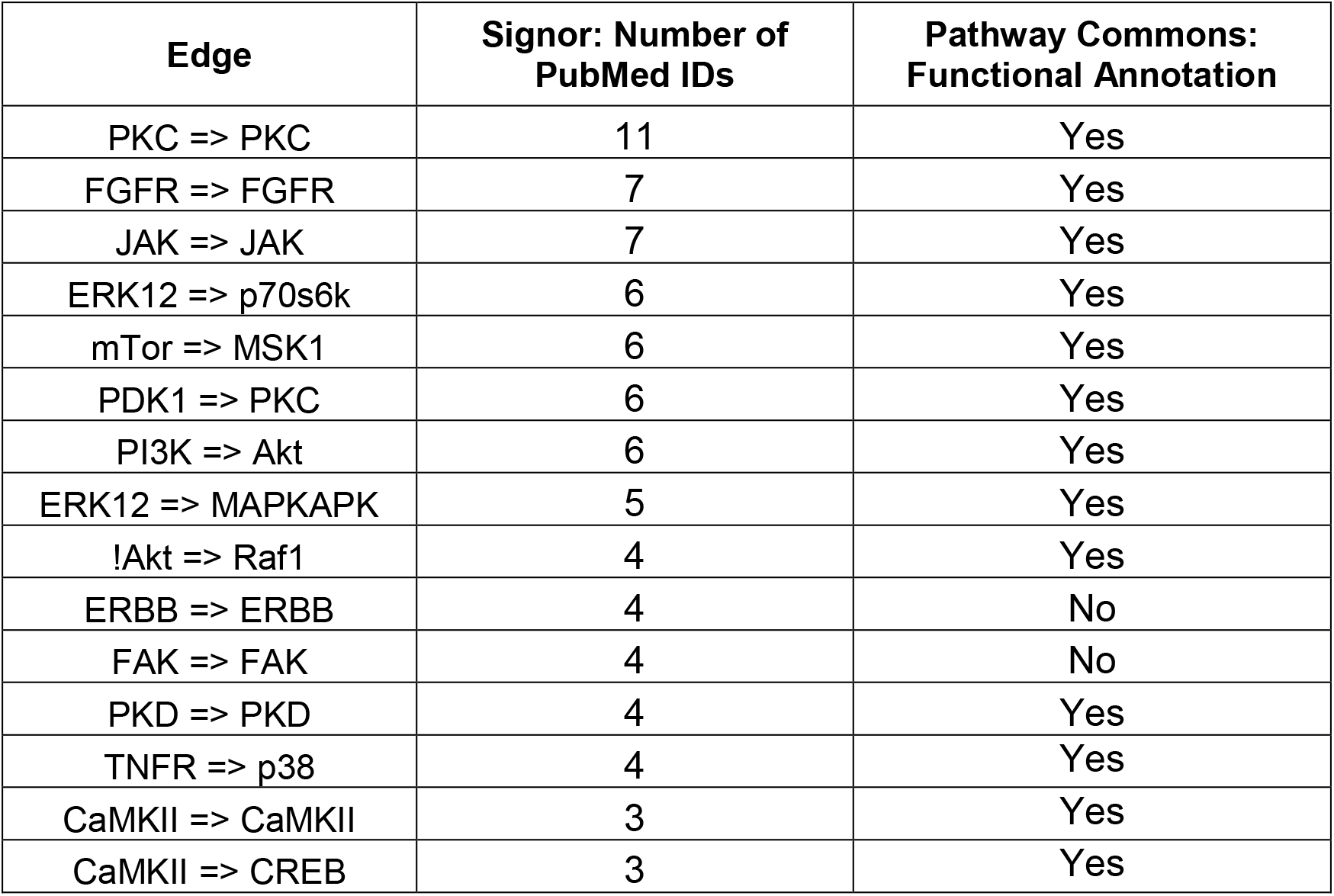
Top 15 candidate edges when ordered by number of Signor PubMed IDs.

| Edge | Signor: Number of PubMed IDs | Pathway Commons: Functional Annotation |
| --- | --- | --- |
| PKC => PKC | 11 | Yes |
| FGFR => FGFR | 7 | Yes |
| JAK => JAK | 7 | Yes |
| ERK12 => p70s6k | 6 | Yes |
| mTor => MSK1 | 6 | Yes |
| PDK1 => PKC | 6 | Yes |
| PI3K => Akt | 6 | Yes |
| ERK12 => MAPKAPK | 5 | Yes |
| !Akt => Raf1 | 4 | Yes |
| ERBB => ERBB | 4 | No |
| FAK => FAK | 4 | No |
| PKD => PKD | 4 | Yes |
| TNFR => p38 | 4 | Yes |
| CaMKII => CaMKII | 3 | Yes |
| CaMKII => CREB | 3 | Yes |

### Integration of candidate edges into logic-based predictions of cardiomyocyte hypertrophy signaling

How would incorporation of new edges obtained from Signor affect the predictions of the logic-based differential equation model of the hypertrophy signaling network? To answer this question, we expanded the hypertrophy signaling network model by incorporating the candidate edges in **Table 4.** At first, candidate edges were added individually, evaluating the performance of the network against a battery of 450 experiments from the prior literature (15). Out of the fifteen candidate edges added to the network, only three individually improved the validation of the model (**Figure 6a**). These were PKC autophosphorylation, ERBB autophosphorylation, and CaMKII phosphorylation of CREB.

**Figure 6.**
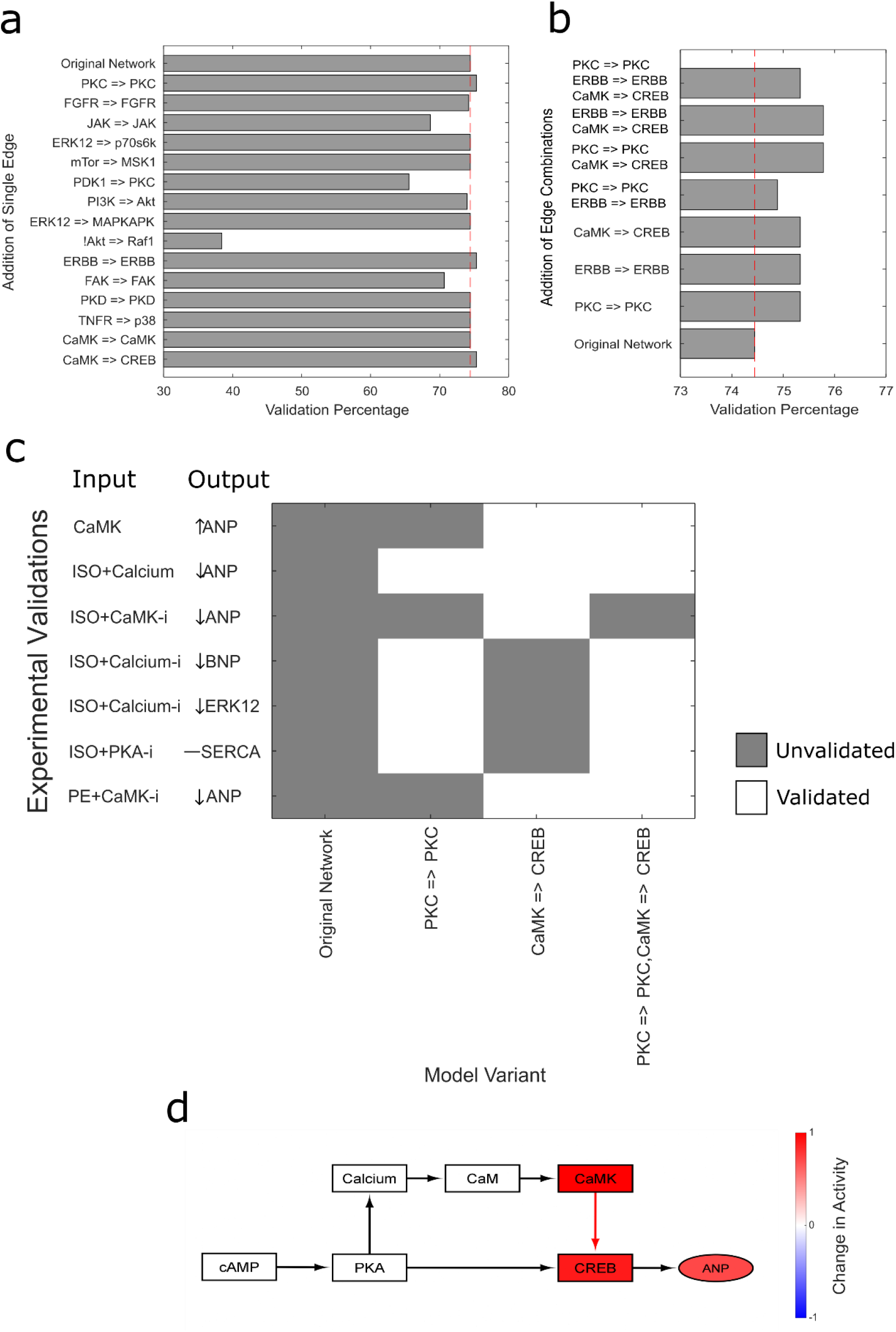
Incorporation of new edges from Signor improves validation of a logic-based hypertrophy network model. Variations in the hypertrophy network model incorporating the top candidate edges from **Table 4** were validated against a battery of 450 experiments from the literature (15). a) Individual edges added to the mode and the impact on model validation. b) Edge pairs that impact model validation. c) Examination of the specific experimental validations that are improved by adding single or paired edges. Mechanistic subnetwork analysis shows how expanding the network with CaMKII=>CREB (red reaction) enables correct prediction of the effect of CaMKII overexpression on ANP activity.

Different combinations of these three edges in the network resulted in varying model performance. One of the best performing combinations was the edge pair of PKC=>PKC and CaMKII=>CREB (**Figure 6b**). The addition of this optimal edge combination to the network resulted in the improved prediction of seven experimental measurements from the literature (**Figure 6c**). A network simulation was performed to visualize the role of the added edges in the improved signaling behavior. To identify the network nodes involved in the overexpression of CaMKII, we performed a virtual knockdown screen and predicted network-wide effects of CaMKII. By overlapping nodes that are sensitive to CaMKII and nodes whose knockdown affects ANP, we identified a mechanistic subnetwork using a previously described method (16). This mechanistic subnetwork highlights the necessity of the CaMKII=>CREB connection (14) to account for the downstream increase of ANP signaling (**Figure 6d**). These results demonstrate that this informed database-driven expansion of the logic-based hypertrophy network improves model predictions.

## Conclusion

Protein interaction databases provide a valuable resource for experimental and computational research, yet the fidelity of these resources requires careful attention. While protein-interaction networks have been benchmarked against focused experiments, they have not been carefully validated for their utility in developing and expanding predictive network models. In this study, we use three large-scale manually curated network models of cardiac hypertrophy, mechano-signaling, and fibrosis to benchmark five protein interaction databases. Overall, we found that roughly 70% of edges in the manually curated networks could be recalled from the top performing Pathway Commons database, and that Signor and Omnipath had particularly high recall for directed interactions given their relatively small size. Network visualization indicated that while Pathway Commons and Signor have strong coverage of central conserved pathways, cardiac-specific reactions and transcriptional regulation had less coverage. To test the utility of protein interaction databases in network model expansion, we ranked candidate edges among proteins in the network by the number of Pubmed references in Signor. This approach identified CaMKII phosphorylation of CREB and PKC autophosphorylation as important for the ability to accurately predict ANP phosphorylation.

While these benchmarking studies indicate that most edges of our network models can be recovered from existing protein interaction databases, this study also reveals several limitations of these databases specific to constructing predictive network models. The most obvious need is that of more interactions incorporated into protein interaction databases. Pathway Commons is the largest database considered here and had the highest recovery rate. However, the quality of curation is also highly important. Signor has a greater degree of manual curation, and indeed while it has only 3.8% as many directed interactions in Pathway Commons, Signor recovered 51% and 45% as many interactions from the mechano-signaling and fibroblast networks respectively. Further, visualization of database annotations on the hypertrophy network topology showed that Pathway Commons and Signor annotate somewhat different regions of the network. Yet gaps remained, particularly for cardiac-specific interaction, which points to the continued need for manual curation for tissue-specific network models.

Network visualization also revealed gaps in coverage of transcriptional regulation in these databases. While this study focused on cell signaling, future studies could use manually curated network models of gene regulation to benchmark databases that focus on transcriptional regulation, such as Chea3 (17). In addition to manually curated resources, there are now a wide range of omic data that could systematically integrate with predictive network models. In particular, data that examine a specific transcriptional regulator and its impact on global gene regulation. For example, CHIP-seq identifies genome-wide binding of particular transcription factors. Rogers et al. expanded fibroblast network model with gene expression data using transcription factor enrichment (18), which was based on GRNBoost2 (19). Another area needing greater coverage and inclusion into models is post-transcriptional regulation, such as the transcription factors (20) and targets (21) of microRNAs.

This study focused on benchmarking protein interaction databases using three logic-based network models of signaling networks. The comparison of these three networks was performed because they were manually curated in a consistent manner, and all three models have been highly validated by experiments. This enabled direct comparisons, which revealed consistent recovery of interactions across hypertrophy, mechano-signaling, and fibrosis networks. While the logic-based formalism of these models was only used for Figure 6, future studies should examine whether network models of other biological systems and with other modeling formalisms have similar recovery rates from protein interaction databases.

An important alternative to manual literature curation and use of protein interaction databases is the use of network inference algorithms, which are typically based on experimental data of the network state (22). The approach described here could be applied to benchmark network inference methods using manually curated network models. Because these network models enable comprehensive simulations, they would enable systematic study of the advantages and disadvantages of various inference methods. This complements previous approaches such as the DREAM competition that validated inference using experimental data not used in inference (23). Some network algorithms even combine prior knowledge from interaction databases with network state data to improve inference (24), and this could be extended to include networks from manually curated models. While powerful, due to limited experimental data the accuracy of network inference methods is typically low, which necessitates the continued use of manual literature curation.

In conclusion, this benchmarking study revealed significant utility in the ability of protein interaction databases for recovering and also predicting new edges for predictive network models. This demonstrates a promising approach for systematically expanding manually curated network models and reveals new insights into cardiac hypertrophy signaling.

## Methods

### Comparing network model topology to protein interaction databases

In benchmarking a curated network model, all possible pairwise gene product interactions corresponding to a given edge were used to represent a single interaction. For example, node A is denoted as activating node C in rule 3 of the toy network (**Figure** 1). The C node represents gene C, and, thus, two pairwise gene product interactions are used to benchmark this edge against databases. In scoring this interaction against a database, each of the gene product interaction combinations is compared to the list of interactions in a database. If either of the combinations is present, the edge is scored as being present in the database. Across an entire network, the performance of a database was determined by calculating the fraction of network interactions represented in a given database. If an interaction included a small molecule or denoted a connection with cellular phenotype, it was excluded from benchmarking.

Because some of the database interactions represent directed links while others are ambiguous with regards to directionality, a separate benchmarking score was computed for the portion of each database that encoded undirected or directed interactions. Interactions not denoted as being directional were checked for forward and reverse matches with network edges. After the fraction of network interactions present was calculated for each database used, the information was merged across database to evaluate for any additive benefits when benchmarking.

### Determining annotations for network edges

The possible genes corresponding to each node in a given network were used to measure Pathway Commons interaction annotations for edges in the curated network model. A given edge was determined to have a functional or physical annotation if any of the pairwise gene product interactions corresponding to that edge had a functional or physical annotation. If an edge had functional and physical interactions, it was classified as having both, while an edge with neither functional nor physical annotations was classified as having no annotations. The PubMed IDs documented in Pathway Commons for all of the pairwise gene product interactions in a network edge was used to generate a list of unique PubMed IDs supporting that network edge. The same process was used to count the number of directed and undirected annotations in Signor as well as to generate a list of Signor PubMed IDs. Edges that included small molecules or denoted connections with cellular phenotype could not be included in benchmarking and were excluded from any benchmarking.

### Expanding the logic-based differential equation model of hypertrophy signaling with new database-driven edges

Candidate network edges identified from protein databases were used to expand the network model. Edges were added individually to create model variations. These variations were used to compare simulations against 450 known experimental outcomes (15), as performed previously. If the validation of the model variations were at least as good as the validation of the original network model then those corresponding edges were considered high performing edges. The high performing edges were then used in combination and compared against the original network using the same validation criteria. The best performing combination of edges was then used to expand the original network model.

Computer code for network benchmarking and running network validation simulations is freely available at https://github.com/saucermanlab/NetworkBenchmarking.

## Acknowledgements

This study was supported by grants from the National Health Institute (R01HL162925, R01HL160665, R01HL137755, T32HL007284) and the University of Virginia Pinn Scholar Award.

